# Asymmetries in foveal vision

**DOI:** 10.1101/2024.12.20.629715

**Authors:** Samantha K. Jenks, Marisa Carrasco, Martina Poletti

**Affiliations:** Department of Brain and Cognitive Sciences, University of Rochester; Department of Neuroscience, University of Rochester; Center for Visual Science, University of Rochester; Department of Psychology, New York University; Center for Neural Science, New York University

## Abstract

Visual perception is characterized by known asymmetries in the visual field; human’s visual sensitivity is higher along the horizontal than the vertical meridian, and along the lower than the upper vertical meridian. These asymmetries decrease with decreasing eccentricity from the periphery to the center of gaze, suggesting that they may be absent in the 1-deg foveola, the retinal region used to explore scenes at high-resolution. Using high-precision eyetracking and gaze-contingent display, allowing for accurate control over the stimulated foveolar location despite the continuous eye motion at fixation, we investigated fine visual discrimination at different isoeccentric locations across the foveola and parafovea. Although the tested foveolar locations were only 0.3 deg away from the center of gaze, we show that, similar to more eccentric locations, humans are more sensitive to stimuli presented along the horizontal than the vertical meridian. Whereas the magnitude of this asymmetry is reduced in the foveola, the magnitude of the vertical meridian asymmetry is comparable but, interestingly, reversed: objects presented slightly above the center of gaze are more easily discerned than when presented at the same eccentricity below the center of gaze. Therefore, far from being uniform, as often assumed, foveolar vision is characterized by perceptual asymmetries. Further, these asymmetries differ not only in magnitude but also in direction compared to those present just ∼4deg away from the center of gaze, resulting in overall different foveal and extrafoveal perceptual fields.

## Introduction

It is well established that perception across the visual field is not uniform. As eccentricity increases, visual acuity ^1,2,3^, contrast sensitivity ^4,5^, object recognition ^6^, and cortical magnification –the amount of cortical surface area corresponding to one degree of visual angle ^7,8,9^– decline. Several factors contribute to this decline, including an increase in retinal cone ^10,11,12^ and retinal ganglion cell ^13,14^ spacing, as well as an increase in the population receptive field (pRF) size in the visual cortex ^15,16^. It is, however, less known that vision is not uniform at isoeccentric locations (polar angle) along a given eccentricity (see ^17^ for a review). In particular, sensitivity differs at isoeccentric locations at the same eccentricity. This phenomenon is referred to as visual asymmetries (*e.g.,* ^18,19,20,21,22,23,24,25,26,27^).

There are two well-studied visual asymmetries in the extrafoveal visual field: the *horizontal-vertical meridian asymmetry*, characterized by greater sensitivity along the horizontal than the vertical meridian, and the *vertical meridian asymmetry*, characterized by greater sensitivity along the lower than the upper vertical meridian (*e.g.*, ^18,19,20,21,22,23,24,25,26,27^). These asymmetries have been assessed in many basic visual tasks, such as contrast sensitivity ^18,19,21,23,26^, texture segmentation ^28^, and acuity ^29,30,31^. Interestingly, the extent of these asymmetries increases with higher stimulus spatial frequency ^18,26,29^ and with increasing eccentricity from the center of gaze ^18,24,32^. It has also been shown that asymmetry is the strongest along the meridian, but gradually diminishes with angular distance from the meridian ^23,24,29^. These changes create a continuous field of gradually varying sensitivity, referred to as the performance field (*e.g.*, ^33,18,19^).

These visual asymmetries have also been reported in more complex tasks such as letter identification task ^33^ visually guided pointing ^34^, motion discrimination ^35^, numerosity ^36^, perceived size ^37^, and visual short-term memory ^38^. Furthermore, visual asymmetries persist even when spatial attention–both exogenous ^18,19,28,39,40^ and endogenous ^41,42^– or temporal attention ^43^ is engaged. Therefore, visual asymmetries are pervasive and shape multiple aspects of visual processing.

These visual asymmetries have been extensively studied in the extrafoveal visual field, yet it is unknown whether these asymmetries extend to the foveola. The foveola receives input from the central 1-degree of the visual field ^44^. This region is responsible for high-resolution vision; it is defined by being devoid of capillaries and rods, and it is characterized by the highest cone density and spatial resolution. Therefore, the foveola is of paramount importance in many everyday tasks such as reading, driving, and discriminating stimuli from a distance. Further, although the foveola covers less than 0.01% of the visual field, its input is over-represented by ≈ 800 times in the primary visual cortex ^45^. Yet, foveolar vision is often assumed to be characterized by uniformly high sensitivity and is generally treated as a single homogeneous region.

Based on evidence that the magnitude of visual asymmetries decreases at near eccentricities ^18,24,46^ and that spotlight sensitivity for isoeccentric locations in the foveolar field is uniform ^47^, we may expect no asymmetries in visual discrimination within the central 1-degree of the visual field, where acuity, under normal viewing conditions, is primarily limited by uniform optics ^48,49,50^. However, evidence showing that fine spatial vision already starts to decline across the central fovea ^51,52^, and that retinal cone density along the vertical meridian of the foveola declines with eccentricity more pronouncedly than along the horizontal meridian ^10,14,12^ would suggest otherwise.

Assessing visual asymmetries at the foveolar scale requires precise localization of the line of sight, with arcminute level accuracy, to present small stimuli at predefined eccentricities from the preferred locus of fixation, a capability that exceeds what commercial video-eyetrackers can achieve ^53^. To overcome these issues and investigate fine visual discrimination at isoeccentric locations in the foveola, we relied on a custom-made high-precision Dual Purkinjie Image eyetracker ^54^ coupled with a gaze contingent display system ^55^. Together, these systems enable the recording of eye movements with high precision and provide a more accurate localization of the line of sight ^54,56^.

## Results

Visual asymmetries are defined as differences in visual perception (*e.g.*, visual sensitivity and acuity) at different polar angles along a given eccentricity. Whereas asymmetries in the parafovea and perifovea have been extensively studied, it is not yet known whether asymmetries are present in the 1 deg central fovea. To investigate this issue, fine spatial discrimination was tested at eight isoeccentric locations in the foveola (4 cardinal and 4 intercardinal), ≈20 arcminutes away from the preferred locus of fixation (Foveolar condition, Fig. 1*A*–*B*). As a comparison, performance was also assessed at the corresponding locations in the parafovea at 4.5 deg eccentricity (Parafoveal condition). Observers were asked to maintain fixation on a central marker while discriminating the orientation of a tiny bar (tilted ± 45 deg) that was briefly presented at one of the tested locations (Fig. 1*A*). To maintain comparable task difficulty between the foveal and parafoveal conditions, stimulus contrast was adjusted to yield an overall performance ≈70% of correct responses across the 8 tested locations in each condition (t(7) = −0.61, p = 0.56, paired two-tailed t-test; BF = 2.54 for the null hypothesis; Fig. 1*C*).

**Figure 1:**
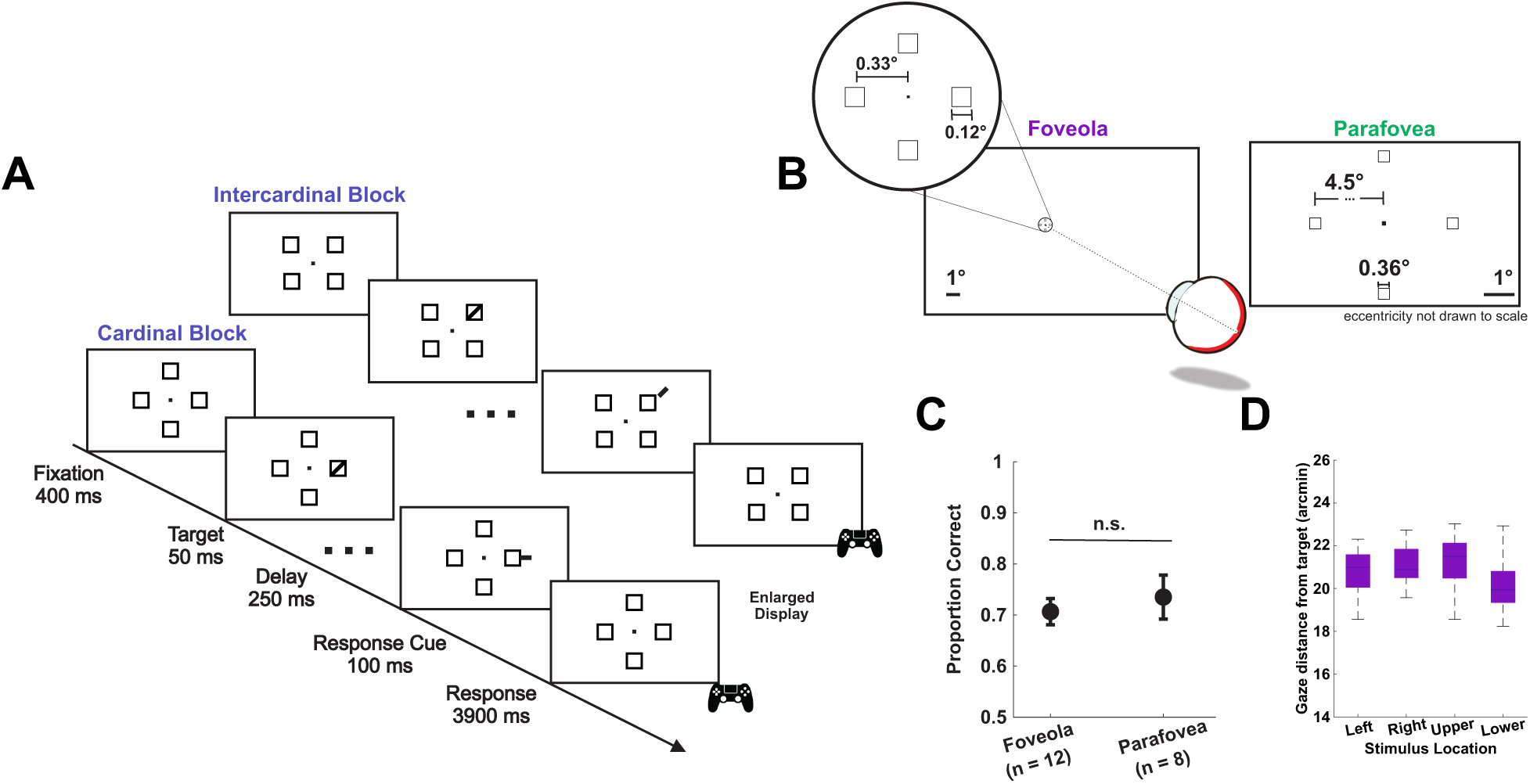
Experimental protocol. **A-** Subjects fixated monocularly (left eye patched) on a central fixation marker while stimuli were presented. The target, a small bar tilted ± 45 deg, appeared briefly (50ms) at one of 4 possible locations and subjects were asked to report its orientation once the response cue was presented. A total of eight locations were tested. Cardinal and intercardinal locations were tested in separate blocks. **B-** Stimuli spatial arrangement and dimensions in the foveola (left; n = 12) and parafovea (right; n = 8) conditions. In the latter condition, stimuli were magnified according to the cortical magnification factor^57^. **C-** In a preliminary session, the stimulus contrast was determined for the foveola and the parafovea conditions separately, so that overall performance across cardinal and intercardinal locations yielded ≈70% correct responses. Error bars are 95% confidence intervals. **D-** The target was 20 arcmin from the center of the display. The box and whisker plot shows that the average gaze distance from the target was approximately 20 arcminutes for different target locations in the foveola condition.

Testing fine spatial vision within the foveola is challenging because it is difficult to present and maintain stimuli at the desired location, only arcminutes away from the preferred locus of fixation. During fixation, eye movements continually shift the retinal projections of objects across the foveola, even during brief fixation periods ^58,59^. Here, we used a high-precision eyetracker ^54^ and a gaze contingent display ^55^ to more accurately localize the line of sight and to either present stimuli at a fixed eccentricity using retinal stabilization, or to post-hoc select only trials in which gaze position was maintained within a circular region of 10 arcminutes in radius around the center of the display (10% ± 7% of trials in the foveola were removed post-hoc for gaze off center). Figure 1*D* shows that in the latter case, the average target distance from the preferred locus of fixation remained approximately at the desired eccentricity of 20’ ± 3’, and it was comparable across all tested locations (N=10, F(7,9) = 1.93, p = 0.08, one-way ANOVA; BF = 1.56 for the null hypothesis). Retinal stabilization was used only for two subjects who showed larger fixational instability (see Methods for details).

Despite the stimuli being presented at isoeccentric locations, performance was not uniform across the eight tested locations both in the parafovea (F(7,7) = 12.21, p *<* 0.0001, one-way ANOVA; BF *>* 100) and in the foveola (F(7,11) = 4.95, p = 0.0001, one-way ANOVA; BF *>* 100). Consistent with previous literature (*e.g.*, ^33,18,19,24,37,26,29,60,36^), in the parafoveal condition, we found the typical horizontal-vertical meridian asymmetry in performance; subjects were on average 27% ± 12% better at discriminating stimuli along the horizontal than the vertical meridian (Figure 2*A*–*B*; horizontal = 91% ± 4%; vertical = 64% ± 3%; two-tailed paired t-test: t(7) = 6.37, p = 0.0004; BF = 91.22; Cohen’s d = 3.44). Interestingly, when stimuli were presented foveally, a similar pattern was found (Figure 2*A*; horizontal: 74% ± 4%; vertical: 68% ± 3%; two-tailed paired t-test: t(11) = 2.71, p = 0.02; BF = 3.36; Cohen’s d = 0.94). This asymmetry was present for all individual subjects in the parafovea and for most subjects in the foveola (Figure 2*B*). However, the magnitude of this asymmetry was 4.4 times larger in the parafovea than in the central fovea (Figure 2*C*). These results are consistent with the findings that asymmetries increase with eccentricity ^18,24,46^.

**Figure 2:**
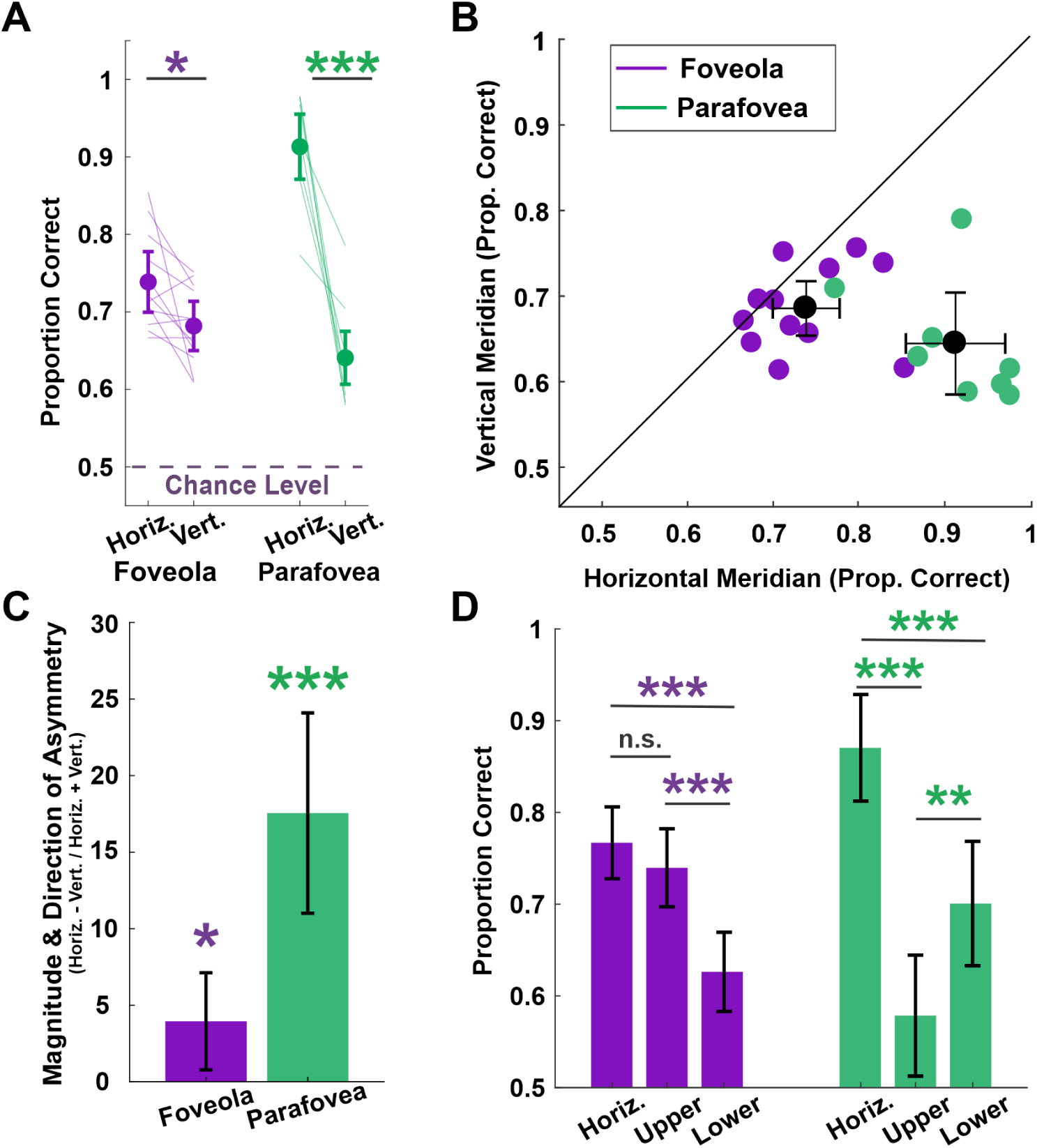
Horizontal-Vertical Meridian Asymmetry. **A and B-** Average performance along the horizontal (pooled left and right locations) and vertical meridian (pooled top and bottom locations) across subjects for the foveola and the parafovea. Lines in **A** and colored dots in **B** represent individual observers. In **A**, asterisks denote a statistically significant difference (*P*<*0.05, **P ≤ 0.01, ***P ≤ 0.001, paired t-test p = 0.02 foveola, p = 0.0004 parafovea). **C-** Magnitude and direction of the horizontal-vertical meridian asymmetry. Asterisks indicate a statistically significant difference from zero (one sample t-test p = 0.02 foveola, p = 0.0004 parafovea). **D-** Average performance along the horizontal meridian compared to the upper and lower vertical meridian performance for the foveola and the parafovea. Asterisks denote statistically significant differences (one-way ANOVA, p = 0.0001 foveola, p *<* 0.0001 parafovea). All Error bars are 95% confidence intervals.

In the parafoveal condition, both locations along the vertical meridian showed a significant drop in performance compared to those along the horizontal meridian (F(2,7) = 37.33, p *<* 0.0001; BF = 5.51; horizontal: 91% ± 4%; lower vertical: 70% ± 7%; upper vertical: 58% ± 7%). However, in the foveola condition, the horizontal-vertical meridian asymmetry was mainly the result of a drop in performance in the lower vertical meridian (F(2,11) = 13.29, p = 0.0002; BF = 106.35; horizontal: 74% ± 4%; lower vertical: 63% ± 4%; upper vertical:74% ± 7%; Figure 2*D*).

There was no statistically significant difference in reaction times for stimuli presented along the vertical vs horizontal meridian in the foveola (Z = −1.2990, p = 0.1939, Wilcoxon rank-sum test), and in the parafovea reaction times were faster for stimuli presented along the horizontal meridian (horizontal: 367ms ± 125ms; vertical: 524ms ± 180ms; Z = −2.15, p = 0.031, Wilcoxon rank-sum test), indicating that the reported asymmetries in visual discrimination are not due to a speed accuracy trade-off.

Besides the horizontal-vertical asymmetry, differences in visual perception are also reported along the vertical meridian. Previous research has consistently shown that in the parafovea and perifovea sensitivity is higher in the lower than the upper vertical meridian (*e.g.*, ^23,18,19,26,29,60,61^). Our findings in the parafovea are in line with the literature (Fig. 3*A* – *B*; lower: 70% ± 7% upper: 58% ± 7%; two-tailed paired t-test: t(7) = −4.90, p = 0.002; BF = 26.21; Cohen’s d = 1.35). In the central fovea, we reported an asymmetry of comparable magnitude (fovea: 11% ± 9% and parafovea: 12% ± 7%), indicating that not all asymmetries are attenuated in this region. Remarkably, the pattern of asymmetry reversed in the foveola (Fig. 3*C*); visual discrimination was better in the upper than the lower vertical meridian (upper: 73% ± 4% vs. lower: 63% ± 4%; two-tailed paired t-test: t(11) = 4.32, p = 0.001; BF = 34.07; Cohen’s d = 1.56). When comparing reaction times in the lower and upper vertical meridians for the foveola and parafovea, there was no statistically significant difference (foveola: Z = −1.18, p = 0.24; parafovea: Z = 0.9977, p = 0.32, Wilcoxon rank-sum test), indicating that the results were not due to a speed-accuracy trade off.

**Figure 3:**
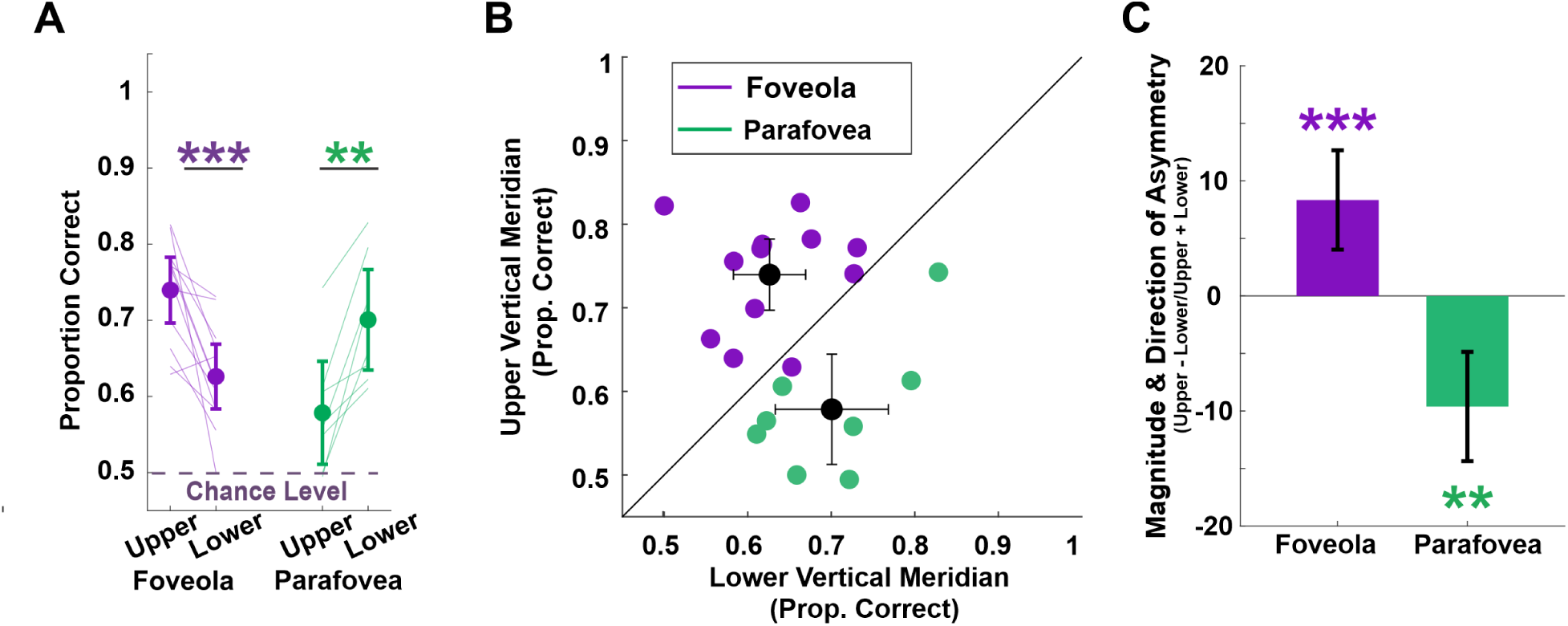
Vertical Meridian Asymmetry. Conventions are the same as in Fig. 2. **A and B-** Average performance along the upper and lower vertical meridian across subjects. In **A**, asterisks denote a statistically significant difference (paired t-test, p = 0.001 foveola, p = 0.002 parafovea). **C-** Magnitude and direction of the vertical meridian asymmetry. Asterisks indicate a statistically significant difference from zero (one sample t-test, p = 0.001 foveola, p = 0.002 parafovea). All error bars are 95% confidence intervals.

To examine the visual performance fields with finer resolution, visual discrimination was also tested at intercardinal locations. As reported in some studies (*e.g.*, ^18,23^), the most noticeable feature of the parafovea and perifovea performance field was the enhancement of performance along the horizontal meridian, giving the performance field an oblong shape (Fig. 4*A*). The performance field in the central fovea, in contrast, is less oblong and it is mainly characterized by an upper vertical meridian enhancement (Fig. 4*A*). Performance between the upper and lower intercardinal locations were comparable for the foveola upper: 73% ± 4%, lower: 68% ± 9%; two tailed paired t-test: t(11) = 1.62, p = 0.13; BF = 1.25 for the null hypothesis; Cohen’s D = 0.51) and the parafovea (upp r: 65% ± 8%, lower: 73% ± 13%; two tailed paired t-test: t(7) = −2.00, p = 0.085; BF = 0.78 for the null hypothesis; Cohen’s D = 0.65). This result is consistent with findings in the parafovea and perifovea ^18,19,23,24,41,29^.

**Figure 4:**
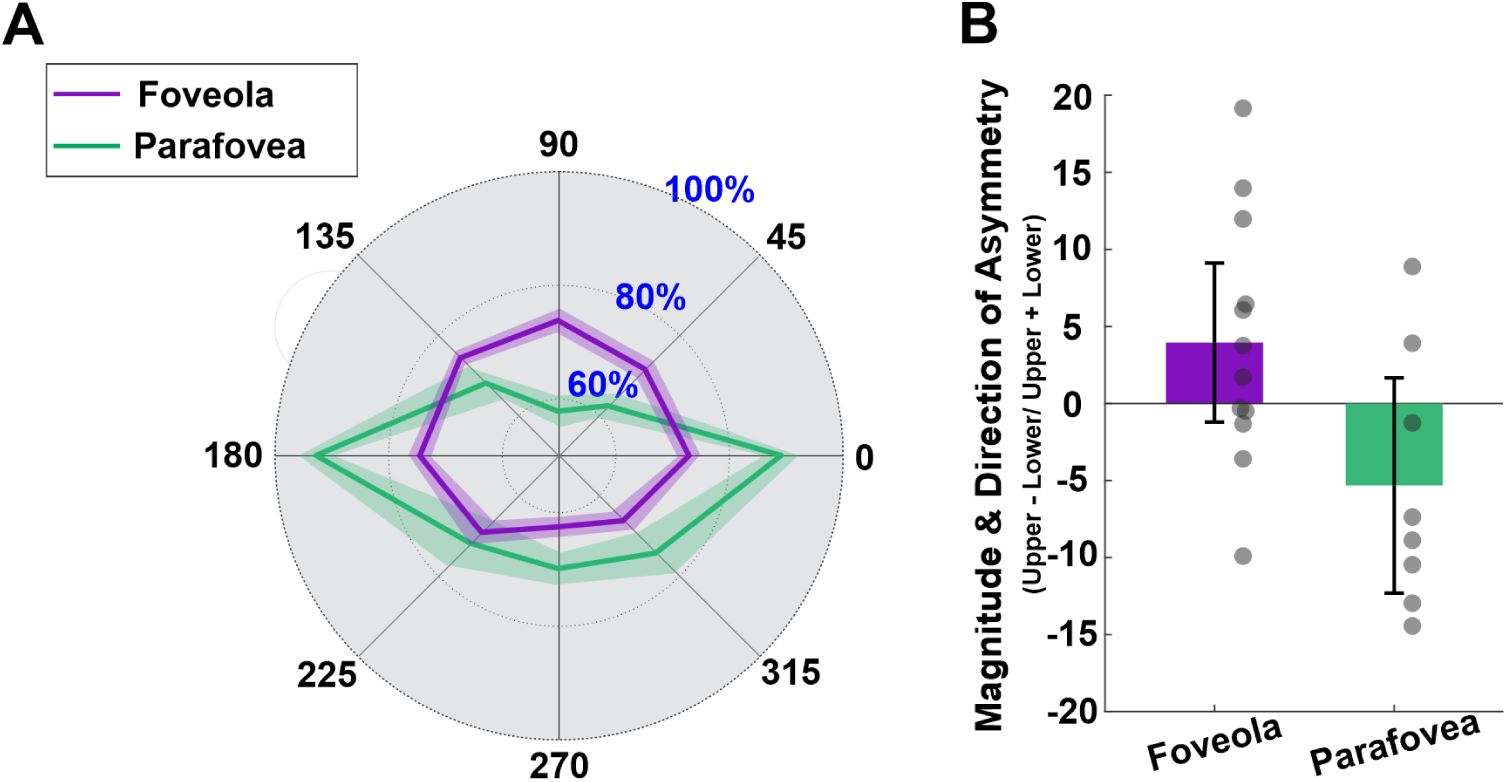
Performance fields in the foveola and parafovea. **A-** The visual performance field in the foveola (purple) and parafovea (green). Each angle represents one tested location. Numbers along the radial direction represent percent correct, and the shaded regions represent SEM. The lines connect the average performance across subjects for the different locations tested. **B-** Magnitude and direction of the asymmetry between the upper intercardinal locations (locations at 45 and 135 deg) and the lower intercardinal locations (locations at 225 and 315 deg). Filled circles represent individual observers. Error bars are 95% confidence intervals.

## Discussion

It is typical to characterize extrafoveal vision at different eccentricities rather than as a whole, as many functions gradually decrease as stimulus information is presented more peripherally ^62,63,64^ (see ^65,66^, for reviews). However, examining foveolar vision as a whole is common practice. Whereas this may seem justified by the fact that the central fovea covers only a tiny (*<*0.01%) portion of the visual field, it is at odds with the fact that the foveola is vastly overrepresented in the primary visual cortex, which dedicates 8% of its surface area to the processing of the visual input coming from this region ^45^. Moreover, cone density ^11,12,14^ and fine visual discrimination ^51,52^ have been shown to decline just a few arcminutes away from the preferred retinal locus of fixation. Therefore, foveolar vision is less homogeneous than usually assumed.

Here, we show that foveolar vision not only changes with eccentricity but also varies around the polar angles of the visual field projecting onto this region, as is the case extrafoveally (see ^17^ for a review). Specifically, we found that discriminating stimuli located degrees below, compared to above, the center of gaze is easier, whereas discriminating stimuli just a few arcminutes below versus above the center of gaze proves more challenging. This foveal asymmetry, although comparable in magnitude, is opposite in direction to the corresponding asymmetry in the extrafoveal visual field. In general, the overall shape of the parafoveal performance field is oblong along the horizontal meridian, whereas the foveolar performance field is primarily enhanced along the upper vertical meridian.

The horizontal-vertical meridian asymmetry has been reported in parafovea and perifovea studies (*e.g.*, ^18,19,20,21,22,23,24,25,26,27^). Notably, this parafoveal asymmetry is more prominent in this study than in many previous studies. This difference may be due to differences in stimulus parameters –stimuli used in previous work are larger and often limited to specific spatial frequencies– and the presence of the placeholders, which could have been perceived as distractors –as asymmetries become more pronounced with distractors ^18,41^. Ultimately, these results show that not only is the foveola characterized by visual asymmetries, but also that these asymmetries are unique in the central most 1 deg of the visual field.

As discussed in the Introduction, extrafoveal asymmetries are pervasive and have been shown across various visual tasks and stimulus parameters, raising the question of where along the visual processing pipeline these asymmetries emerge. The first factor that could influence the visual asymmetries are the optics of the eye. Optical aberrations are not symmetric along the visual field meridians ^67^, and horizontal/vertical astigmatism and coma vary as a function of polar angle ^68^. Specifically, the magnitude of vertical coma gradually decreases from the inferior to super retina (across −18 to 18 deg), resulting in more optical distortion from vertical coma in the upper compared to the lower vertical meridian ^68^. In addition to optical factors, at the anatomical level, the extrafoveal retina is characterized by higher cone ^69,70,10,71^ and retinal ganglion cell ^13^ density along the horizontal than the vertical meridian. Whereas retinal cone density contributes towards the horizontal-vertical meridian asymmetry, no difference in cone density has been reported along the vertical meridian. However, the density of midget-retinal ganglion cells (mRGCs) is higher along the superior retina, which projects to the lower visual field, compared to the inferior retina ^13^. Moreover, in the mid-periphery, the convergence of cones to mRGCs is minimal along the horizontal meridian but is the greatest along the inferior retina ^13^, resulting in greater information loss for stimuli in the upper visual field. This difference aligns with behavioral evidence from this and previous studies.

A computational observer model, however, has shown that optical and retinal factors account for about 40% of the behavioral horizontal-vertical asymmetry and about 10% of the behavioral vertical meridian asymmetry, and that they are amplified at later processing stages in the visual cortex ^72,73^. In the primary visual cortex asymmetric differences in BOLD response ^74^, population receptive field size ^75,17^, and cortical surface area ^76,8,77,17^ have been reported. Specifically, for a given eccentricity range, the horizontal meridian can have up to 80% greater cortical surface area than the vertical meridian, and a similar pattern is seen for the lower vs. upper vertical meridian ^15,8,77,17^. Although further investigation is necessary to elucidate the degree to which the visual cortex influences perceptual asymmetries, factors from the retina to the cortex increasingly contribute to the reported visual asymmetries.

What factors influence foveal asymmetries? Unlike in the extrafovea, optical aberrations in the foveola are constant ^48,49,50^. Therefore, we can exclude the influence of optics on foveolar asymmetries. However, retinal factors could play a role in shaping these perceptual asymmetries. Midget retinal ganglion cells (mRGC) have a 1:1 mapping with retinal cone cells ^44^, and thus neither mRGC density nor cone-to-mRGC convergence should impact the pattern of asymmetries. It has been observed that the foveola exhibits similar asymmetries in cone density to those in the extrafoveal retina, with a higher cone density along the horizontal than the vertical meridian ^12,10^. This is consistent with our findings that fine visual discrimination is better along the horizontal vs. the vertical meridian. However, like in extrafovea, the foveola shows no differences in cone density along the vertical meridian ^12^. Therefore, it would seem that cone density alone cannot account for the visual asymmetries observed in the foveola.

Importantly, changes in cone density with eccentricity are generally defined with respect to the peak cone density (PCD) location. However, this location does not coincide with the retinal projection of the center of gaze. The center of gaze on the retina is considered as the preferred retinal locus of fixation (PRL), which can be quantitatively defined as the median retinal location of a stimulus during fixation. Notably, there is an offset between the PRL and the point of highest cone density ^78,12,79,80,81,82,83,47^. Specifically, the PRL is shifted nasally and superiorly from the cone density centroid (CDC)–the centroid of the region with highest cone density–by approximately 5 arcminutes ^12^. Although this is a small shift, it introduces systematic asymmetries in cone density between the superior vs. inferior retina. Therefore, if eccentricity is defined with respect to PRL rather than from the CDC location, cone density is higher along the inferior retina, corresponding to the upper visual field, than along the superior retina. Although the overall performance was comparable at the intercardinal locations due to relatively large individual variability, this difference in cone density below vs. above the preferred retinal locus could potentially contribute to the vertical meridian asymmetry observed in the foveola. It is likely that subsequent stages of processing in the visual cortex further contribute to the foveolar perceptual asymmetries. Yet currently, unlike in extrafoveal vision, nothing is known about possible cortical asymmetries in the processing of foveolar visual input.

Visual sensitivity differences at isoeccentric locations along different directions may serve functional purposes in vision. A horizontal enhancement could be beneficial for socialization with other people at the same height range as faces are commonly located along the horizontal meridian. This asymmetry is also advantageous for reading, both in the fovea and parafovea, as humans can extract words 12-15 letters ahead of the fixated word along the horizontal meridian ^84,85^. Interestingly, the horizontal-vertical meridian asymmetry is present in both children and adults ^86^, suggesting that it may develop early in life or even be present from birth. However, the vertical meridian asymmetry is absent in children, both in behavior ^86^ and cortical surface area ^17^; behaviorally, it emerges in adolescence ^87^. A lower vertical visual field enhancement might be beneficial as the objects that we typically interact with are usually located below our center of gaze ^88^; however, they are no longer present at intercardinal locations ^18,19,23,24,41,89^.

Whereas there are no clear functional advantages for the vertical meridian asymmetry at the foveal scale, this asymmetry may be related to oculomotor factors. Specifically, it has been shown that upward saccades greater (but not smaller ^60,30,27,61,90^) than 10 degrees tend to undershoot, whereas downward saccades of the same amplitude tend to overshoot the target stimuli ^91^. As a result, the foveated target falls above the center of gaze every time a large vertical saccade, either upward or downward, is performed. Therefore, better fine visual discrimination in the upper foveal visual field may be advantageous in these circumstances.

What is the effect of attention on the foveal asymmetries? Covert attention enhances visual performance at the attended location (see ^92^ for a review). In the extrafovea, exogenous attention enhances sensitivity uniformly across isoeccentric locations, thereby preserving the magnitude and pattern of the visual asymmetries ^18,19,39,40^. Interestingly, despite generally being characterized by a more flexible deployment ^93,89^ endogenous attention was also found to increases the performance field uniformly ^41,42^. Therefore, covert spatial attention allocates comparable resources at all attended locations regardless of whether they are along the horizontal or vertical meridian in the extrafovea. It has been established that attention can be selectively deployed within the foveola, both endogenously ^94^ and exogenously ^95^. Yet, unlike in the extrafovea, it is not known whether fine scale attention has an effect on foveal asymmetries.

In conclusion, using a high-precision eyetracker to investigate visual performance at selected locations across the 1-degree foveola, this study revealed that foveolar vision is characterized by visual asymmetries in fine visual discrimination. Whereas the magnitude of the horizontal-vertical asymmetry was attenuated compared to the corresponding parafoveal asymmetry, the magnitude of the asymmetry along the vertical meridian (i.e., upper vs lower) was comparable in both conditions. Remarkably, however, the direction of the vertical asymmetry was reversed in the fovea indicating that distinct mechanisms are at play at this scale. This difference may be the result of fixation behavior leading to changes in cone sampling above vs below the fixated location, given the slight offset between the peak cone density location on the retina and the preferred retinal locus, or it may arise from a different cortical representation of foveal input at the level of V1 and beyond. Importantly, these results further emphasize the need to consider foveal vision not as a uniform entity, but as one characterized by significant non-uniformities that shape perception of fine detail.

## Methods

### Subjects

The experiment included 13 participants (11 na ive), including one of the authors, aged 18 or older (mean 24 ± 4.54 years; 8 female) with normal or corrected to normal vision. One subject was removed from the analysis due to performance being at chance level in all conditions tested. Eight of the twelve subjects participated in both conditions (foveola, n=12 and parafovea n=8). The University of Rochester Institutional Review Boards approved the experiment. All study participants provided consent prior to the study.

### Apparatus

Stimuli were displayed on an LCD monitor with a refresh rate of 200 Hz and a spatial resolution of 1920 × 1080 pixels (Acer Predator XB272). The task was performed monocularly with the right eye while the left eye was patched. To prevent head movements, a unique dental-imprint bite bar and a headrest were used. The right eye’s movements were recorded at 340Hz using a Basler-acA2000-340KM-NIR camera. EyeRIS, a custom-developed system that allows flexible gaze contingent display control, was used to render the stimuli ^55^. This system acquires eye movement signals from a high precision digital Dual Purkinje Image eyetracker ^54^, processes them in real time, and updates the stimulus on the display based on the desired combination of estimated oculomotor variables.

### System Calibration

Before the start of each block, subjects align their unique dental-imprint bite bar to the center of monitor, by looking through a set of two pinholes. Once aligned, the subject undergoes two step system calibrations. First, subjects undergo a standard automatic calibration procedure using a 3×3 grid of points. Points were 1.32 degrees apart from each other in the horizontal and vertical directions. The mapping obtained through the automatic calibration was then refined using a custom-made manual calibration procedure ^56,51^. This dual-step calibration allows a more accurate localization of gaze position than standard single-step procedures, improving 2D localization of the line of sight by approximately a factor of three on each axis ^51,56^. During the experiment, the manual calibration procedure was then repeated for the central fixation marker before each trial to compensate for possible small head movements introducing errors in gaze localization.

### Experimental Protocol

After calibration, subjects started the trial by pressing a button. The stimulus array consisted of four 7×7 arcminute squares at 0.33 deg (foveola condition) or 22×22 arcminute squares at 4.5 deg (parafovea condition) away from the central fixation square (5×5 arcmin). These placeholders could be either located along the cardinal directions or along four intercardinal directions. Cardinal and intercardinal directions were tested in separate blocks for a total of eight tested locations. While subjects maintained fixation on a central marker, a small bar (2×7 arcmin or 6×20 arcmin) titled ± 45 deg was briefly presented for 50 ms at one of four possible locations. The parafoveal stimuli were magnified according to the cortical magnification factor at that eccentricity ^57^, similarly to the way is calculated in prior studies ^96,97,46,98^. Subjects performed a 2AFC orientation discrimination task. Stimuli contrast was adjusted using the method of constant stimuli during a preliminary session to yield an overall performance of 70% correct responses across the eight locations tested. Stimuli contrast was then maintained at this level throughout the following experimental sessions. After 250 ms from target offset, a green response cue (2×7armin or 6×20 bar) appeared for 100ms. Subjects had four seconds to report the orientation of the target stimulus using a joypad.

### Analysis of oculomotor data

Only trials with optimal, uninterrupted tracking and free from saccades and microsaccades during the period of interest (50ms before target onset to 50ms after target offset) were selected for data analysis. When retinal stabilization was used trials characterized with drift amplitudes larger than 100 arcmin were removed to eliminate instances in which subjects attempted to chase the stabilized stimulus (0.2% of trials for subject 1 and 1.2% for subject 5). When stimuli were viewed without retinal stabilization in the foveola condition, trials in which the gaze was ≥ 10 arcmin away from the central fixation marker during the period of interest (50ms before target onset to 50ms after target offset) were discarded to ensure that stimuli were presented at the desired eccentricity (see Methods) (on average 10% ± 7% of trials were removed for this reason). This criterion was more relaxed in the Parafoveal condition, as the stimuli were farther away from the center of gaze. Trials in which gaze was ≥ 30 arcmin away from the central fixation marker during the period of interest were discarded (on average 2%± 1% of trials).

### Analysis of behavioral data and statistics

Besides characterizing performance as percent of correct responses we also calculated performance as d-prime. Hit and false alarm rates were corrected for response bias (Stanislaw and Todorov, 1999).Overall, we found the results based on the d-prime performance in agreement with those based on percent correct reported in the main text. Both conditions, showed a significant horizontal-vertical meridian asymmetry; participants were on average 58% ± 24% better at discriminating stimuli along the horizontal than the vertical meridian in the parafovea (horizontal = 3 d’ ±1 d’; vertical = 0.75 d’ ± 0.4 d’; two-tailed paired t-test: t(7) = 5.43, p = 0.0001; BF = 41.96; Cohen’s d = 2.86), and 15% ± 18% and better in the foveola (horizontal: 1.34 d’ ± 0.41 d’; vertical: 0.97 d’ ± 0.28 d’; two-tailed paired t-test: t(11) = 2.69, p = 0.021; BF = 3.28; Cohen’s d = 0.98). Additionally, our findings show that the sensitivity in the lower vertical meridian was higher than in the upper vertical meridian (lower: 1.12 d’ ± 0.5 d’ upper: 0.42 d’ ± 0.42 d’; two-tailed paired t-test: t(7) = −4.94, p = 0.002; BF = 27.05; Cohen’s d = 1.35). Again, this pattern was reversed in the foveolar condition; the upper vertical meridian was more sensitive than the lower vertical meridian (upper: 1.36d’ ± 0.42 vs. lower: 0.67d’ ± 0.38d’; two-tailed paired t-test: t(11) = 4.46, p = 0.001; BF = 41.15; Cohen’s d = 1.58.)

All analyses were performed in MATLAB. ANOVAs, post-hoc multiple comparison tests, paired t-tests and Cohen’s D calculations were performed using MATLAB’s statistical toolbox. To quantify the magnitude of the effects we calculated the BayesFactor for ANOVA’s and t-tests using the Bayesfactor MATLAB toolbox (https://zenodo.org/badge/latestdoi/162604707)

## Acknowledgements

This work was funded by NIH R01 EY029788-01 to MP, NIH training grant T32EY007125 to SJ, EY001319 and NIH NEI Grant R01-EY-027401 to MC.

